# Uncovering cognitive-motor adaptation to body visualization in immersive technologies through their behavioral and electrophysiological correlates

**DOI:** 10.64898/2026.09.02.748913

**Authors:** Elpidio Attoh-Mensah, Arnaud Boujut, Tristan Delaval, Clément Naveilhan, Stephen Ramanoël, Anaick Perrochon

## Abstract

Immersive technologies, encompassed within the x-reality (XR) framework spanning the reality-virtuality continuum, are emerging as tools to facilitate behavioral adaptation through interaction with controlled environments. Yet, how XR features shape users’ behavior remains poorly understood, particularly when a coordination between cognitive and motor demands is required. Among these features, body visualization (*i.e*., how the user’s body is rendered within the environment), is especially relevant to XR usages in rehabilitation and skill training, since body-related feedback is known to strongly influence movement adaptation under cognitive–motor interference. Still, whether and how different body visualization scenarios shape movement adaptation and what are the underlying neural processes, remains unclear.

To address these questions, thirty healthy young adults performed a stepping task concurrently with an updating n-back task under three body visualization scenarios (real-body, no-body, and a knee-position cue) and four levels of dual-tasks while we recorded high-density mobile electroencephalography. The absence of body visualization impaired motor performance, reducing stepping accuracy and increasing omissions. Remarkably, a knee-position cue alone restored performance to real-body levels, without improving embodiment. This behavioral benefit co-occurred with neural modulations reflecting enhanced conflict monitoring and adjusted motor preparation, evidenced by changes in the frontocentral N450 component and sensorimotor alpha-and beta-band activity, respectively. These findings indicate that minimal body-related visual cues, rather than full-body visualization, can support cognitive-motor performance in XR, without altering embodiment. They position cueing as a practical design strategy for movement-related XR usages and motivate further investigations of other forms of body-related feedback to optimize cognitive-motor behavior.

## 1. Introduction

Immersive technologies are transforming practices in various domains including industry, research, and healthcare (1). Their description has been first conceptualized under the reality– virtuality continuum also known as extended reality which spans real environments to fully virtual ones (2). Between these two extremes, augmented and mixed realities overlay virtual elements onto the real world and may enable interaction with virtual objects in the physical surroundings (2, 3). However, this foundational concept remains inconsistent across studies because the experiences cannot be fully characterized only by the proportion of real and virtual elements, or by the display of the technology. Recent taxonomy have updated the reality-virtuality continuum by introducing the x-reality (XR) framework where “x” stands for a new form of reality (4). This new XR framework suggests that the user experience would be multidimensional incorporating features of the technology, its content, and their effect on users (4). Technology and content-related features include principally immersivity (*i.e*., possibilities of perceptual and cognitive absorption), interactivity (*i.e*., possibilities of enacted behavior within the environment), and explorability (*i.e*., possibilities of navigation) (3–5). These are complemented by other content-related features which determine how users can perceive the environment as plausible and believable (5). Together, all these features would affect higher-order user experience such as presence, the psychological sense of “being there”, and embodiment which involves body ownership, self-location, as well as agency (6).

Careful optimization of XR features has been widely recommended across various contexts (7, 8), given their critical role in shaping the user experience. For instance, the user’s sense of embodiment directly influences enacted behavior within the environment. Recent meta-analytic evidence (9) suggested that current evaluations of the sense of embodiment within XR largely trace back to the experiment of the rubber hand illusion which demonstrated that seeing a limb being stimulated in synchrony with one’s own hidden limb can induce a sense of ownership over the seen limb (10). Accordingly, the sense of embodiment in XR is principally modulated through body visualization, which influences self-perception and interaction within the environment (11, 12). Interestingly, the effects of body visualization on the sense of embodiment have been attributed to the congruence of available sensorimotor feedback (9, 13–16). This is particularly important, since it may explain why body visualization would affect enacted behavior within the environment. Indeed, previous reports suggested that the absence of body visualization amplifies the impact of disrupted sensorimotor contingencies during upper-body movements in virtual reality (13). Other authors reported that despite having no direct effect on embodiment components, visuo-tactile congruency induced through manipulations of the body visualization altered both body schema and walking behavior (14). These findings are particularly relevant in XR since environments differ in the availability, precision, and congruence of body-related feedback. Indeed, in virtual reality, users typically act without seeing their body or through a virtual body representation, whereas in augmented and mixed realities they may retain visual access to their real body while interacting with virtual elements.

Existing findings support the relevance of body visualization for movement in XR, but the evidence remains fragmented, limiting the ability to select technology that is closely aligned with the intended objectives whenever movement is involved. In virtual reality, evidence supports that the absence of body visualization tends to degrade movement quality compared to partial or full-body visualization, especially in reaching, grasping, and hand-based tasks (15–17). In augmented and mixed realities, synchronizing a virtual body with the real body has been shown to improve performance in gait and hand-reaching tasks (18, 19). However, studies comparing different XR environments are scarcer with inconsistent results. While some authors reported faster execution of finger-pointing movements in augmented reality, involving real body visualization, than in virtual reality with virtual body representation (20, 21), some other reported shorter grasping movement duration and better interaction (22). Several factors may explain these inconsistencies, including differences in the tasks performed across studies and the type of technology used whose features may differentially impact the outcomes. This lack of consistency echoes a research gap recently outlined in perspectives on XR usages suggesting the need for a better understanding of the role of XR features towards the intended goals (23–25). Filling this gap is particularly important when it comes to body visualization and related movement adaptations. Indeed, beyond the limited control on the technology-related features, the few existing studies comparing XR environments relied on relatively simple motor tasks (20, 21) that do not reflect the complexity and variety of movements.

In XR usages, movements are often goal-directed and combined with cognitive demands, resulting in cognitive-motor dual-task situations (1, 26–28). During such tasks, cognitive-motor interference (CMI) can emerge when available attentional resources are insufficient to manage motor and cognitive demands simultaneously (29). As a result, individuals adopt different strategies depending on the situation (*e.g*., prioritizing the motor task, prioritizing the cognitive task, or attempting to balance priority between both) (30). CMI and the associated strategies are highly relevant as they can serve as diagnostic markers (31), targets for rehabilitation or skill training (32, 33), or experimental tools to investigate cognitive-motor adaptations (29, 34). The implementation of these strategies is governed by a combination of both top-down (*e.g*., optimal allocation of attentional and executive resources depending on the context) and bottom-up processes (*e.g*., real-time adaptation based on sensorimotor feedback) (29, 35) whose effectiveness is further impacted by moderating factors such as the functional capacities of participants (36, 37), and characteristics of the tasks (38). In XR usages, body visualization scenarios may constitute an additional moderator by modifying the availability, precision, and congruence of the sensorimotor feedback. When this feedback is reduced or incongruent, the information available to guide action may become less reliable, which could affect CMI strategies. Conversely, richer and more congruent body-related feedback may facilitate adaptation of the movement.

Behavioral measures can provide valuable insights as they enable the assessment of cognitive and motor performance decrements and/or improvements associated with CMI (29). However, these measures alone may provide limited insight into the specific contribution of the strategy *per se*, independently of the aforementioned moderating factors. Indeed, individuals may exhibit similar patterns of cognitive–motor adaptation while relying on different strategies to achieve these outcomes (39). Therefore, beyond behavioral outcomes, it is important to gather additional inputs to better understand the underlying processes of movement adaptation under CMI. At this point, Mobile brain/body imaging is particularly relevant because it allows the recording of neural activity during free movement, thereby capturing the real-time interaction between environmental features, sensorimotor feedback, and cognitive-motor demands (35, 40–42). For instance, mobile electroencephalography (EEG) offers high temporal resolution, and previous research identified markers of cognitive and motor processes during CMI. Studies based on event-related potentials (ERPs) demonstrated that post-event frontocentral negativities in N200 (~200–350 milliseconds [ms]) and N450 (peaking ~300–500 ms) increase in response to the detection and monitoring of attentional conflicts, respectively (43, 44). In addition, centroparietal positivity P300 (~300– 600 ms) has been associated with the dynamic of the allocation of attentional resources to perform the tasks (45, 46). In time-frequency domain, analyses of event-related spectral perturbations (ERSPs) showed an increase in frontal midline theta (4-8Hz) during cognitive control and modulation of beta (13-30Hz) in sensorimotor areas related to motor adaptation (40, 46). Although the value of the integration of EEG-based assessments with XR setups has been outlined recently (42), the potential of this approach to better characterize body-visualization-related cognitive–motor adaptation in XR remains unexplored.

To address this caveat, this within-subject cross-sectional study aimed to determine how different scenarios of body visualization in XR can shape cognitive-motor adaptation under increasing cognitive-motor interference. In our principal hypothesis, the absence of body visualization would impair cognitive-motor adaptation compared with both virtual body and real-body visualization, particularly under higher levels of CMI. Specifically, we expect lower motor and cognitive performance in the absence of body visualization, with stronger decrements as cognitive and/or motor load increase. We further anticipate that virtual-body visualization would support performance comparable to that during real-body visualization, assuming that there was sufficient congruent body-related feedback available to guide the action. At the neural level, the main hypothesis is that during CMI increase, the following figures would arise in ERPs (increase in N200, N450, and decrease in P300) and oscillatory dynamics (increased frontal midline theta and sensorimotor beta modulation). We further anticipated that such neural adaptations would differ between virtual and real-body visualization, even in the absence of behavioral differences, indicating that neural markers may capture adaptations not observable at the behavioral level. Furthermore, since the effects of the body visualization are related to the congruence of sensorimotor feedback, we expected that they would manifest through the detection and monitoring of conflict (N200 and N450) rather than resource allocation (P300). Accordingly, ERSPs would show a modulation of motor preparation (beta band modulation) rather than increased cognitive load (frontal midline theta).

## 2. Materials and methods

### 2.1 Participants and sample size calculation

Young and healthy participants were recruited from students at the University of Limoges and required to be able to perform moderate-intensity physical activity. Any motor impairment or physical condition incompatible with the tasks employed constituted a non-inclusion criterion. We did not include participants who reported a previous cybersickness experience during immersive exposure. The study was approved by the national ethic committee in sport science research (n° IRB 00012476-2024-27-05-315) and conducted in accordance with the declaration of Helsinki.

We computed a *priori* sample size estimation using a Monte Carlo simulation approach based on linear mixed-effects models (LMM) and implemented in Python using the *statsmodels* package. Several datasets were simulated according to the planned experimental design, including three conditions of XR and four levels of dual-task, with repeated measures for each participant. The simulated model included fixed effects for condition, level, and interaction, as well as random intercept for subjects. The primary effect of interest was the condition × level interaction. The interaction effect was anticipated based on a moderate estimation informed by previous studies reporting larger effects in movement accuracy (η^2^p > 0.20) (20, 21). We adopted this conservative estimate to avoid overestimating statistical power given differences in task complexity as previous studies used simple finger-pointing tasks. Variance components were chosen to reflect realistic inter-individual variability and residual noise (σ_intercept = 0.80, σ_slope = 0.20, σ_residual = 0.60). For each sample size (ranging from 10 to 60 participants), 2,000 datasets were simulated and analyzed using the following LMM (*y ~ condition + level + condition × level + (1* | *participant)*). The sample size with 24 participants reached the expected targets of 0.05 of significance and 0.90 statistical power. Anticipating frequent data loss in MoBI experiments, we inflated this number by 25% aiming at 30 participants in the present study.

### 2.2 Tasks and XR environments

To reduce hardware-related variability, we implemented all XR environments in the Meta Quest 3 headset (Meta, USA), allowing both virtual and mixed realities, and tasks were developed in Unity (Unity Technologies, 2021). Three experimental conditions were implemented and designed to differ primarily in the availability and format of body-related feedback: i) real-body visualization ii) visualization of a cue representing knee position and iii) no-body visualization (Figure 1A). In real-body visualization condition, virtual objects of the tasks were displayed in the real world, and users were able to interact with them. In the two other conditions, the experimental room was virtually reconstructed from a photogrammetric capture of the real world, allowing either the addition of a knee-position cue or the removal of body visualization. This ensured that the only difference between conditions was the visual representations of the body or the cue on the knee position. The cue was a virtual representation consisting of a white sphere marking the knee position. This minimal manipulation was implemented to isolate the effect of a subtle modification in body visualization. The motion tracking relied on an inside-out system based on embedded cameras. Controllers were attached at the knee level to capture flexion movements required during the motor task.

**Figure 1.**
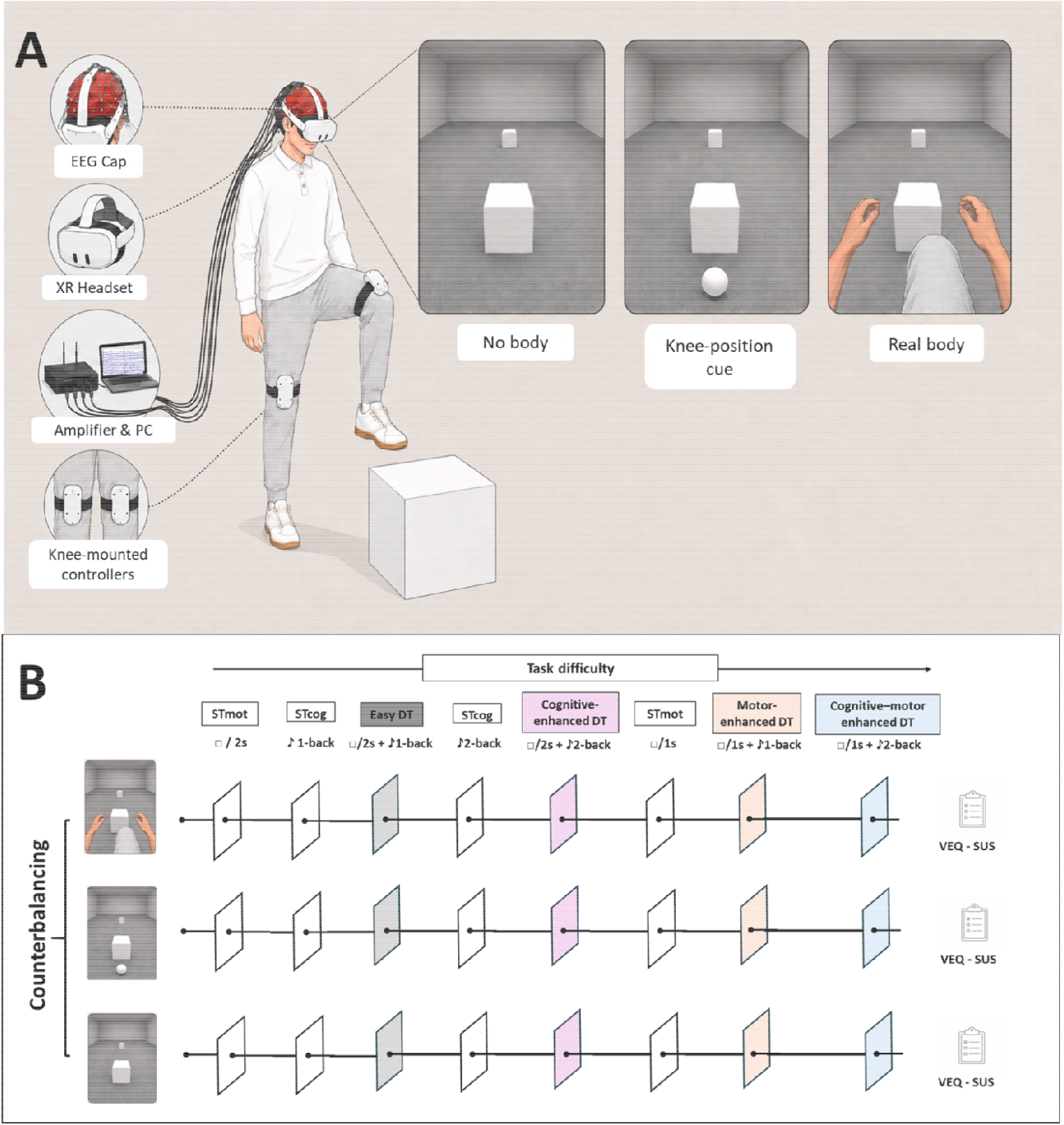
Experimental setup and procedures. **(A)** schematic representation of an equipped participant performing the correct stepping movement. The electroencephalography (EEG) cap is a 64-channel Waveguard Original cap connected to an ANT Neuro eego™ mylab amplifier with 24-bit resolution. The XR Headset was a Meta Quest 3 technology used to implement the scenarios of body visualization. **(B)** Procedures and task completion. Participants completed three counterbalanced experimental conditions based on body visualization scenarios, separated by at least 48 hours to minimize fatigue and carryover effects. After each condition, virtual embodiment questionnaire (VEQ) and slater-ssoh-steed (SUS) were used to investigate embodiment and sense of presence, respectively. Each ~30-min condition comprised 2.5-min blocks of single motor, single cognitive, and various cognitive–motor dual tasks, separated by 1-min rest periods. Dual-task difficulty was manipulated independently through motor load (one obstacle every one or two seconds) and cognitive load (1-/2-back), resulting in four difficulty combinations. Single-task blocks preceded their corresponding dual-tasks to enable the calculation of dual-task costs. Blocks followed a fixed progression to promote gradual adaptation and minimize overwhelming effects.

In the motor task, participants performed stepping movements in response to visual stimuli, consisting of stepping over white cubic blocks appearing on the ground or striking them with the knee when they appeared at a higher position. The stepping task was selected because it involves gait initiation, a key component of cognitive-motor adaptation (31). We conducted an individual calibration phase to adjust the height of the obstacles and stepping movement to each participant’s anthropometric characteristics. Consequently, a correct stepping movement in the task required approximately 90° of hip–knee flexion. Motor load was manipulated by varying stimulus frequency, with an easy stepping level (one obstacle every two seconds) and a hard stepping level (one obstacle per second). The obstacles appeared pseudo-randomly on the left and right sides and on the ground or at knee height. Participants received visual feedback once they successfully cleared an obstacle, which turned green.

In the cognitive task, participants completed a modified version of an auditory n-back task (47), a widely used paradigm to assess working memory and updating processes. Participants heard a sequence of numbers presented every two seconds and for each number, they were required to indicate verbally “yes” if the number was even and “no” if it wasn’t. Two levels of difficulty were implemented. In the easy level (1-back), participants responded based on the immediately preceding stimulus, whereas in the hard level (2-back), responses were based on the stimulus presented two positions earlier. Numbers between 0 to 99 were presented and pseudo-randomized to ensure no repetition (2-2) and ordered series (1-2-3).

### 2.3 Study design and procedure

We counterbalanced the sequence of the three conditions across participants to minimize the effects of a presentation order. We assigned conditions pseudo-randomly to the first and second positions in a way that ensured equal representation across the sample, and the remaining condition was presented last. A minimum interval of 48 hours between conditions was respected to allow washout and reduce fatigue and carryover effects.

During the first session, participants provided anthropometric data and completed the physical activity questionnaire using the short version of the international physical activity questionnaire to assess self-reported active behavior during the previous seven days (48). After each session, participants completed the virtual embodiment questionnaire which assesses body ownership, agency, and perceived body-schema change (49) and the slater-usoh-steed questionnaire which assesses the sense of presence (50).

Each experimental condition lasted approximately 30 minutes and comprised multiple blocks combining varying levels of motor and cognitive load (Figure 1B). These included single motor tasks, single cognitive tasks, and dual-task with different levels of difficulty. Each block lasted 2 minutes and 30 seconds (s), followed by a 1-minute rest period between blocks. The following four blocks of dual-tasks were considered: Easy dual-task (easy stepping + 1-back), cognitive-enhanced dual-task (easy stepping + 2-back), motor-enhanced dual-task (hard stepping + 1-back), and cognitive–motor enhanced dual-task (hard stepping + 2-back). The n-back stimulus presentation consisted of a maximum of 75 trials per block across all conditions. The stepping obstacle presentation included a maximum of 73 trials per block in the easy dual-task and cognitive-enhanced dual-task blocks, and 146 trials per block in the motor-enhanced dual-task and cognitive-motor-enhanced dual-task. Blocks were presented in a fixed order, progressing from easy (single-task) to difficult (dual-task with increasing load levels). We put single tasks just before their corresponding dual-task to be able to use them to compute dual-task cost which reflects CMI. The choice of a fixed order is motivated by prioritizing gradual performance adaptation and limited overwhelming effects over concerns regarding learning effects (30).

### Electroencephalography recordings and preprocessing

We recorded brain activity continuously using a 64-channel Waveguard Original cap equipped with Ag/AgCl electrodes and connected to an ANT Neuro eego™ mylab amplifier with 24-bit resolution. The reference and ground electrodes were positioned at CPz and AFz, respectively, and the sampling rate was 500 Hz. The electrodes’ impedance was maintained below 10 kΩ. EEG recordings were synchronized with behavioral events from stimulus presentation and stepping execution, using time markers via LabStreamingLayer and its dedicated plugin for unity (51), and all data were recorded using the integrated LabRecorder system.

We used MATLAB (R2024b), EEGLAB, and the BeMobil pipeline (52) to conduct the preprocessing offline and in line with previous MoBI protocols (53). Data were first downsampled to 250 Hz, and segments with inactivity, artifacts, or obvious muscle activity were removed manually. We used Zapline-plus to clean spectral noise (*e.g*., line noise, screen refresh), followed by identification of noisy electrodes (on average, 6.5 ± 2.4 per participant), based on channel correlation and maximum broken-time criteria over 10 iterations, which were removed and reconstructed via spherical interpolation. The data were then re-referenced to the common average. We applied a transient high-pass filter at 1.5 Hz before running independent component analysis with the AMICA algorithm (2000 iterations, 10 rejections, 3 SD threshold). We used DipFit and ICLabel in lite mode, for dipole modeling and to classify components, respectively (54). Independent components identified as brain-related (≥ 30% brain probability and < 15% residual variance; (53)) were retained (on average, 13.14 ± 3.7 per participant), and this information was transferred to the unfiltered cleaned data.

### 2.5 Variables analyzed

#### 2.5.1 Self-reported user experience and behavioral data

Self-reported measures included the mean total score on the different questionnaires. Then we analyzed performances within the four levels of dual-task (easy, cognitively enhanced, motor-enhanced, and cognitive–motor enhanced). Motor performance was assessed as the stepping accuracy based on the individual calibration (success = Amplitude of step >= 90° of hip-knee flexion) and the omission (participants did not attempt the task or did not time their step to when the obstacle was presented). Cognitive performance was evaluated through the response accuracy in the n-back task. Subsequently, we calculated dual-task costs across the four levels to quantify the interference between motor and cognitive demands. It was computed using the following formula: [(single-task performance − dual-task performance) × 100 / single-task performance].

#### 2.5.2 ERP and ERSPS extractions

We applied a bandpass filter (0.3–50 Hz) on the continuous preprocessed EEG data and segmented it into epochs (−3s to +3s) time-locked to the stepping execution (time zero corresponding to the moment of clearing the obstacle by hitting it when it is on knee level or avoiding the collision when it is at ground level). Subsequent analyses included exclusively trials with correct stepping execution. We identified artifact-contaminated epochs using a composite quality metric combining maximum epoch amplitude and global deviation from the participant’s average EEG activity. Within each dual-task level, the top 5 % of epochs with the highest artifact scores were rejected. Furthermore, to control for potential effects of unequal trial numbers on phase-based measures, for each participant, we randomly selected an equal number of trials (based on the minimum correct trials in a given dual-task level across body visualization scenarios) within each dual-task level, ensuring balanced trial counts. In each condition, this resulted in an average of 50.17 ± 14.3 epochs per participant and per dual-task level.

In ERPs, we applied a baseline correction using the −200 to 0 ms interval preceding the stepping execution. We averaged the waveforms across electrodes belonging to *a priori* region of interest (ROI) where peak activity during the considered component has been identified. In the frontocentral ROI (Fz, FCz, FC1, and FC2), we investigated N200 and N450 by targeting a window of 160 to 500 ms (41, 43). The P300 component (300 – 600 ms) was further extracted in a centroparietal ROI (Cz, CP1, CP2, C3, C4, and Pz; (45, 55, 56). Time–frequency representations were computed using Morlet wavelet convolution with 3 base cycles and a 0.8 cycle scaling factor. Then, in ERSPs analyses, each trial was first normalized over the full epoch, followed by a re-normalization of the trial averages to the pre-event baseline (−300 to −100 ms) enabling Z-score outputs (40, 57). We focused theta (4–8 Hz) analyses on a frontocentral ROI (Fz, FCz, FC1, and FC2) which covers the frontal-midline and anterior cingulate cortex and is shown to be associated with overall cognitive control (58). The ROI (FCz, FC1, FC2, Cz, C1, C2, C3, and C4) in beta band (13–30 Hz) was defined to encompass the sensorimotor region, with a particular focus on the primary motor cortex, as movement-related beta ERSPs are well characterized within these electrodes (59– 61). Furthermore, we also explored alpha band (8-13 Hz) in the sensorimotor ROI given studies suggesting similar patterns of ERSPs in alpha and beta bands during movement (40, 60, 62).

### 2.6 Statistical Analyses

To investigate the relevance of the manipulation of body visualization, we conducted a one-way repeated-measures ANOVA on self-reported measures of the user experience. Subsequently, we performed behavioral investigations in confirmatory analyses using LMMs on cognitive and motor performances as well as dual-task costs. We considered dual-task levels, body visualization scenarios, and their interaction as fixed effects and participant as random intercept. Observations exceeding three standard deviations from the mean were excluded from the analyses.

Four participants could not be included in EEG analysis due to software issues. Data of the 26 participants included were analyzed using a cluster-based permutation model, which applies ordinary least-squares modelling to continuous EEG while accounting for repeated observations (63). For both ERP and ERSP analyses, the design matrix was constructed to model the factorial structure of the experiment, including body visualization scenarios, dual-task level, and the condition-by-dual-task interaction and participant as random intercept. ERP analyses were performed on time-resolved voltage amplitudes extracted from the mean value of electrodes in the selected ROIs over the time windows of N200 and N450 in frontocentral and P300 in centroparietal ROIs. ERSP analyses included baseline-corrected time-frequency power in the following bands (alpha, beta, theta) and ROIs (frontocentral and sensorimotor), from 800 ms before to 800 ms after the event. We choose this analysis window to minimize overlap in cases of 1s of inter-trial latency which happen in motor-enhanced and cognitive– motor enhanced dual-task levels. The design matrix in both ERP and ERSP was permuted 5,000 times to estimate a null distribution, and main effects and interaction effects were first tested, followed by pairwise comparisons.

Given the continuous nature of the stepping task and the risk of overlap between trials, the pre-event baseline could have been contaminated by the activity of the previous trial due to the general state of the condition. Therefore, we conducted sensitivity analyses on raw amplitudes for ERPs and raw frequency power for ERSP, using the same design matrix as in the principal test by applying an analysis of covariance where baseline values were included as a covariate.

The statistical significance of all tests was set at p < 0.05, and a sequential Bonferroni correction was applied in post-hoc and to control for multiple comparisons. We performed all statistics in SPSS 24.0® (IBM; Armonk, NY, USA) or with built-in MATLAB codes (R2024b). Graphs were also generated with MATLAB. Main effect sizes are represented by partial eta square (η^2^p) and pairwise comparisons effect size by Cohen’s d.

## 3 Results

The 30 participants included had a mean age of 21.9 ± 1.7 and a mean body mass index of 22.9 ± 2.7 kg/m^2^ with 16 women (53.3%) and 14 men (46.7%). The mean score in the international physical activity questionnaire suggests a high level of physical activity in the week preceding the experiment (11181.0 ± 1298.3 in metabolic equivalent).

### 3.1 Self-reported user experience and behavioral data

#### 3.1.1 Validation of experimental manipulation

We identified a main effect of the body visualization in all the sub-sets at the virtual embodiment questionnaire (F_(2, 58)_ = 41.33 to 43.20, *p* = 0.0001, η^2^p = 0.58 to 0.62, Figure 2A-C). Participants reported higher agency and ownership but lower change in body schema during the real-body than both no-body visualization or that of the knee-position cue (*p* = 0.007 to 0.001 and 0.002 to 0.0001; *d* = 1.25 to 1.30 and 1.17 to 1.50, respectively). We observed a similar pattern for the slater-usoh-steed score (F_(2, 58)_ = 7.26, *p* = 0.004, η^2^p = 0.20, Figure 2) with higher presence during real-body than both no-body visualization or that of knee-position cue (*p* = 0.028 and 0.015; *d* = 1.10 and 1.77, respectively). No-body visualization and that of the knee-position cue showed no difference for the two questionnaires (*p* = 0.097 and 0.90, respectively).

**Figure 2.**
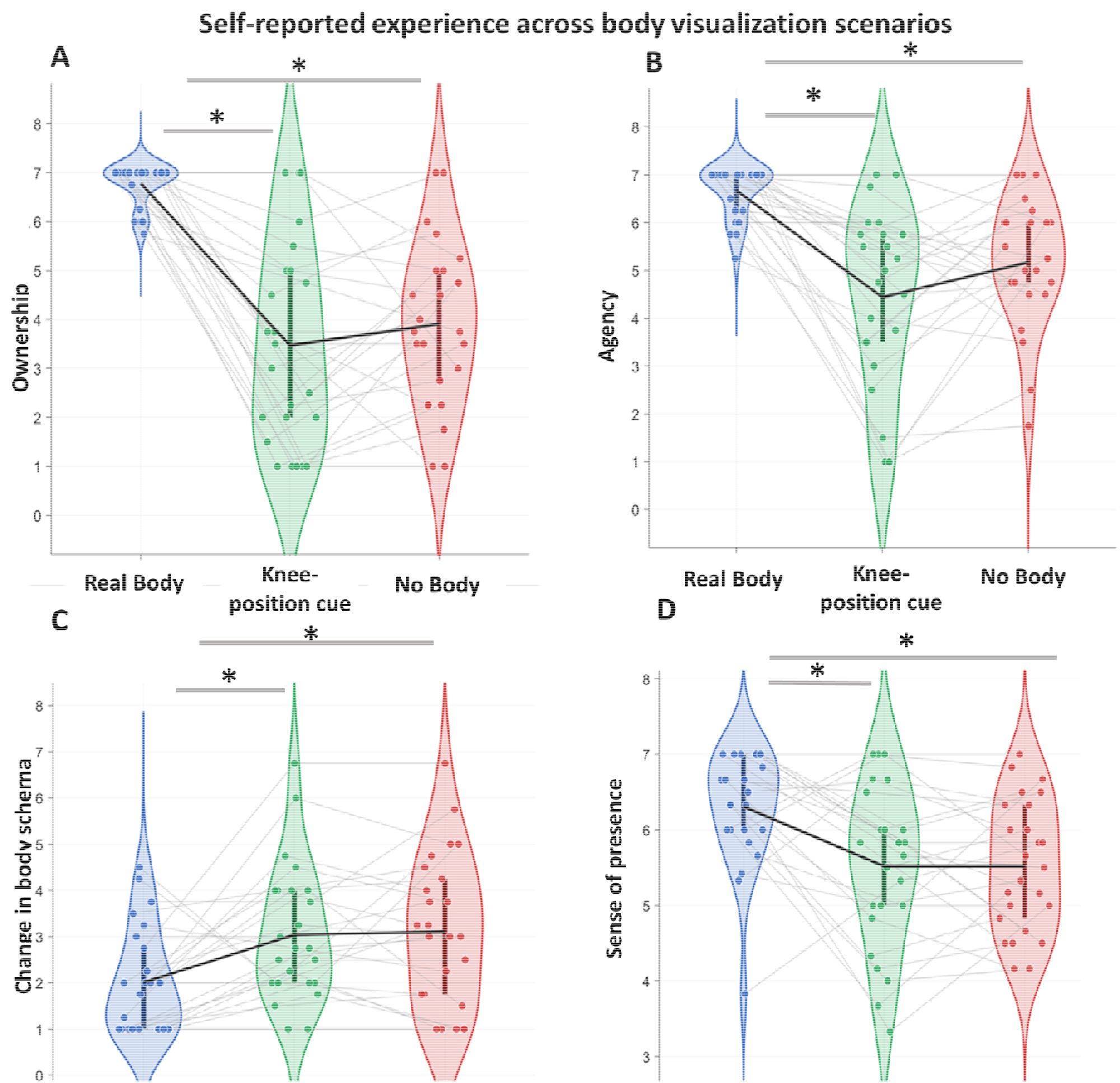
Self-reported experience across body visualization scenarios. **(A-C)** panels show scores on the virtual embodiment questionnaire regarding ownership, agency and change in body schema respectively, while the **(D)** panel shows slater-usoh-steed scores assessing the sense of presence. Distributions are displayed as violin plots with individual data points. Vertical bars within violin plots indicate the interquartile range (25th–75th percentiles). The solid black line across conditions represents the group mean, while gray lines represent individual trajectories. Asterisks indicate significant pairwise comparisons following a significant main effect in the linear mixed-effects model, with sequential Bonferroni correction for multiple comparisons.

#### 3.1.2 Motor performances

LMMs revealed a main effect of the body visualization (F_(2, 318)_ = 4.48, *p* = 0.013, η^2^p = 0.03, Figure 3A) and dual-task levels (F_(3, 318)_ = 9.26, *p* = 0.001, η^2^p = 0.12, Figure 3B) on stepping accuracy. As for stepping omission we identified an effect of body visualization (F_(2, 318)_ = 7.05, p = 0.001, η^2^p = 0.07, Figure 3A) but not for dual-task level (F_(3, 318)_ = 0.69, *p* = 0.560). Stepping accuracy was lower, and stepping omission was higher during no-body than real-body visualization (*p* = 0.037 and 0.001; *d* = 0.35 and 0.46, respectively) and that of the knee-position cue (*p* = 0.040 and 0.002; *d* = 0.30 and 0.37, respectively). No significant difference in accuracy or omission was observed between real-body visualization and that of knee-position cue (*p* = 0.90 and 0.54). As for dual-task levels, stepping accuracy was surprisingly lower in the easy and cognitive-enhanced levels compared to the motor-enhanced and cognitive-motor-enhanced levels. Significant differences were found between the easy and motor-enhanced (*p* = 0.002, *d* = 0.52), easy and cognitive-motor-enhanced (*p* = 0.0001, *d* = 0.64), cognitive-enhanced and motor-enhanced (*p* = 0.031, *d* = 0.42), and cognitive-enhanced and cognitive-motor-enhanced (*p* = 0.003, *d* = 0.52). No interaction effect was identified either for the stepping accuracy (F_(6, 318)_ = 0.81, *p* = 0.55) nor for the stepping omission (F_(6, 318)_ = 1.42, *p* = 0.21). For dual-task costs, there was neither body visualization (F_(2, 318)_ = 0.26, *p* = 0.7) nor dual-task levels (F_(3, 318)_ = 2.85, *p* = 0.40) nor interaction effects (F_(6, 318)_ = 0.49, *p* = 0.812). Mean raw performances are displayed in supplementary materials (Table S1).

**Figure 3.**
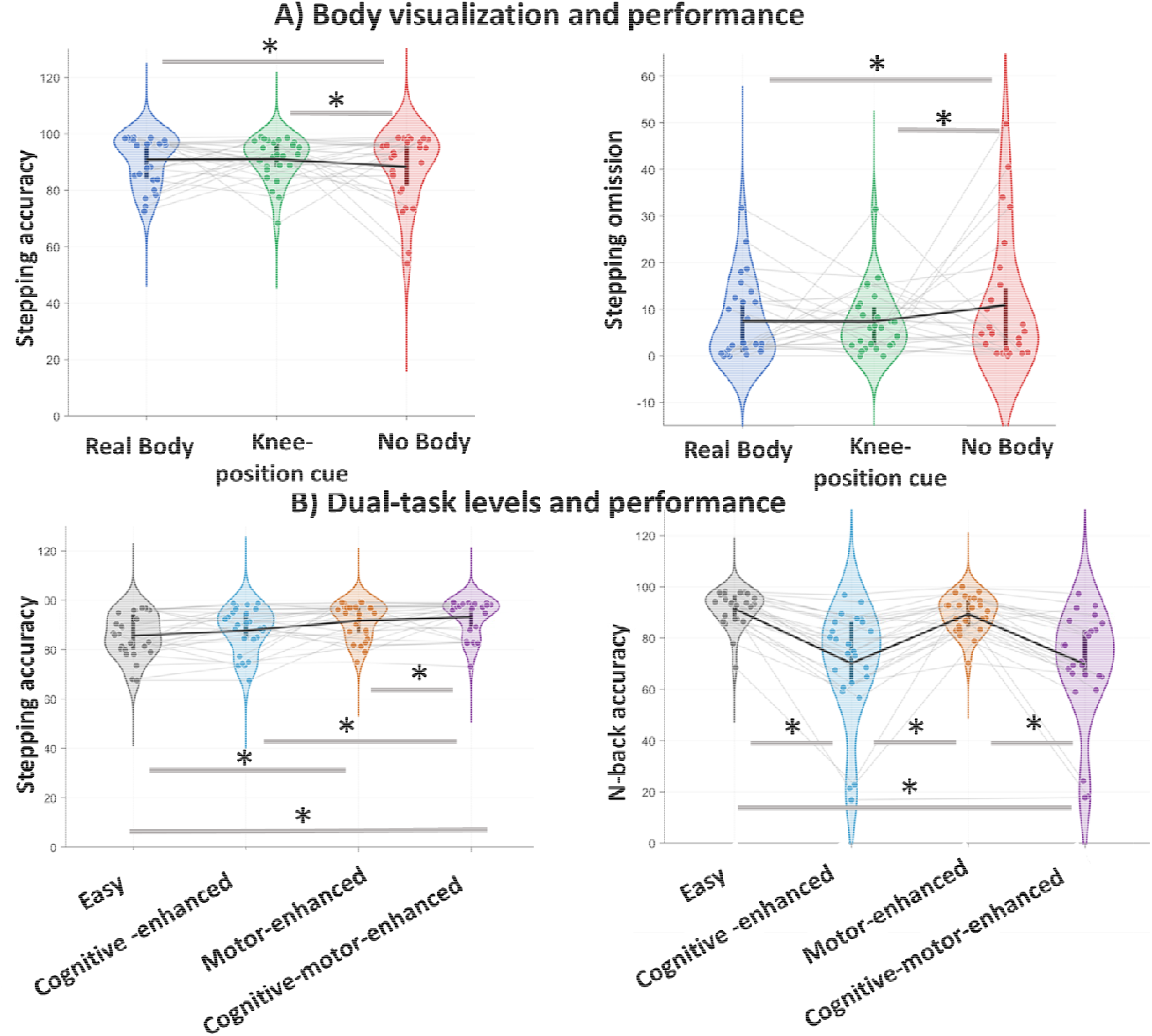
Effects of (A) body visualization and (B) dual-task level on behavioral performances. Distributions are displayed as violin plots with individual data points. Vertical bars within violin plots indicate the interquartile range (25th–75th percentiles). The solid black line across conditions represents the group mean, while gray lines represent individual trajectories. Asterisks indicate significant pairwise comparisons following a significant main effect in the linear mixed-effects model, with sequential Bonferroni correction for multiple comparisons. Sensitivity analyses confirmed these differences while controlling for apparent outliers.

#### 3.1.3 Cognitive performances

We found a main effect of dual-task level for n-back accuracy (F_(3, 318)_ = 16.68, *p* = 0.0001, η^2^□ = 0.42, Figure 3B). This effect was driven by the increase of the cognitive load between two levels: easy *vs* cognitive-enhanced or cognitive-motor-enhanced (*p* = 0.001 and 0.0001; *d* = 1.24 and 1.26, respectively), and motor-enhanced *vs* cognitive-enhanced and cognitive-motor-enhanced (*p* = 0.001 and 0.0001; *d* = and 1.14, respectively). There was no main effect of the body visualization (F_(3, 318)_ = 0.97, p = 0.37) on n-back accuracy. For the dual-task cost on n-back-accuracy also, there was neither body visualization (F_(2, 318)_ = 1.5, *p* = 0.22) nor dual-task levels (F_(3, 318)_ = 3.07, *p* = 0.30) nor interaction effects (F_(6, 318)_ = 1.78, *p* = 0.105).

### 3.2 Electroencephalography data

Details on the exact time windows and duration of the effects on ERPs and ERSPs are reported in the supplementary materials (Table S2 to Table S5).

#### 3.2.1 ERPs

We found a significant main effect of dual-task level for the N450 (F_3,263_ = 6.95, *p* = 0.0001, η^2^□ = 0.06, Figure 4A) and P300 components (F_3,263_ = 5.06, *p* = 0.002, η^2^□ = 0.04, Figure 4A), but not for the N200. The easy and cognitive-enhanced levels elicited comparable responses with both levels displaying significantly smaller N450 and larger P300 amplitudes than the motor-enhanced and cognitive-motor-enhanced levels (*p* = 0.04 to 0.002; *d* = 0.30 to 0.43). No main effect of body visualization was observed for any of the components. However, we found an interaction for the N450 component (F_6,263_ = 2.42, *p* = 0.031, η^2^□ = 0.04, Figure 4B) which was primarily driven by the motor-enhanced and cognitive-motor-enhanced levels with larger N450 amplitudes during the real-body than no-body visualization (*p* = 0.041 to 0.079; *d* = 0.31 to 0.33) while the visualization of knee-position cue was intermediary. The effects of dual-task level (F_3,263_ = 7.71, *p* = 0.0001, η^2^□ = 0.08) and interaction (F_6,263_ = 2.76, *p* = 0.047, η^2^□ = 0.03) on N450 were robust to the sensitivity analyses. This was not the case for the dual-task level effect on P300 due to baseline differences.

**Figure 4.**
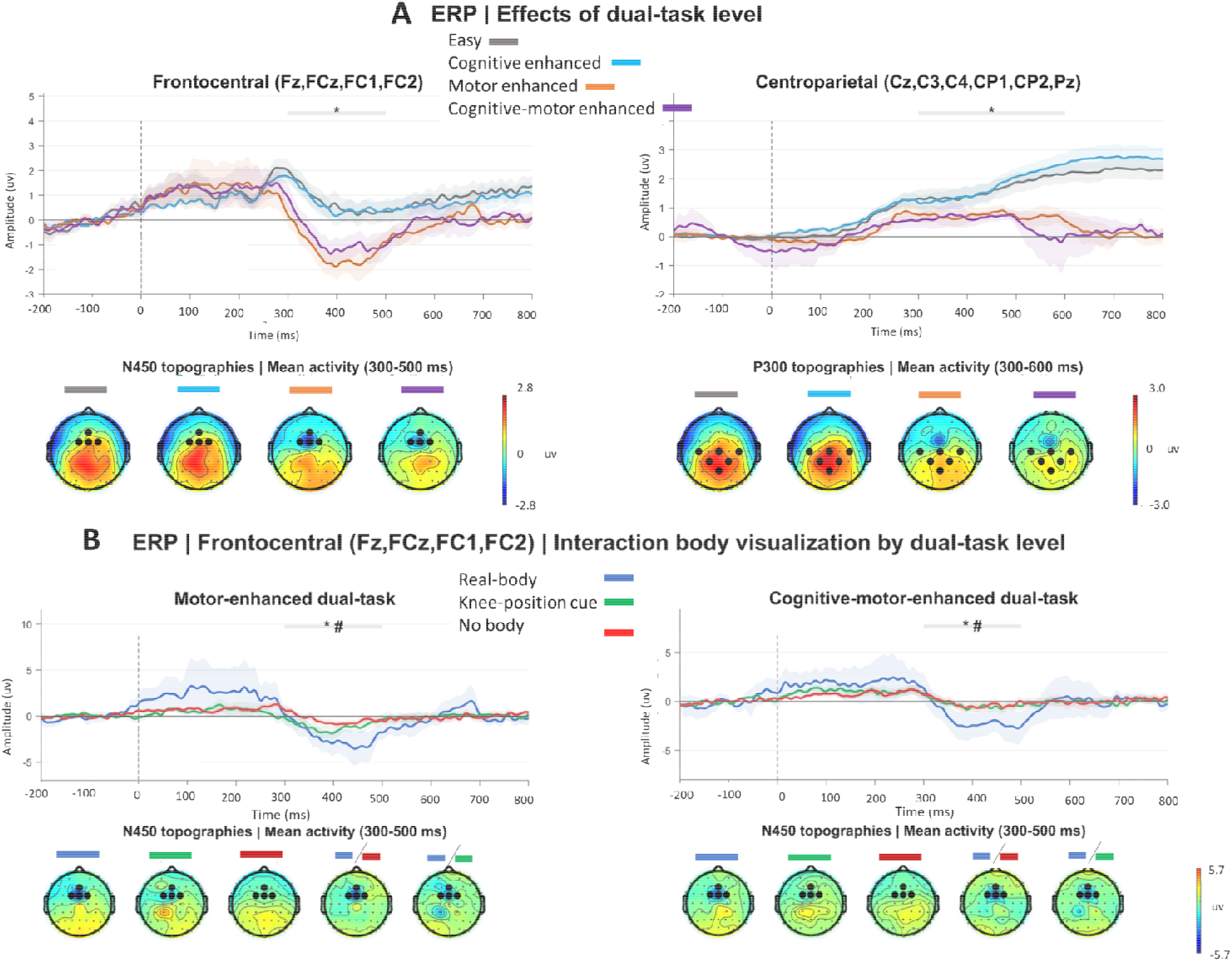
Event-related potential (ERP) across (A) dual-task levels and (B) the interaction between body visualization and dual-task level. ERP panels show time-resolved amplitudes with the corresponding standard error of the mean (SEM). **Time zero represents the moment of clearing the obstacles during the stepping execution**. Asterisks indicate time windows with significant (*p* < 0.05) pairwise differences identified following cluster-based permutation testing, while hashtags indicate statistical trends (*p* < 0.059). **(A)** For both the N450 and P300 components, the easy and cognitive-enhanced dual-task conditions showed comparable amplitudes, both differing from the motor-enhanced and cognitive–motor-enhanced conditions. **(B)** Real-body visualization differed from the no-body visualization condition, while the visualization of knee-position cue, showed an intermediate response. Scalp topographies illustrate the spatial distribution of the components, with small dots indicating all recorded electrodes and larger bold dots highlighting the electrodes included in the region of interest. Only the N450 effects (both the main effect of dual-task level and its interaction with body visualization) remained robust in sensitivity analyses of raw amplitudes when baseline activity was included as a covariate.

#### 3.2.2 ERSPs

Regarding body visualization we found pre-event effects in theta (F_3,263_ = 5.64, *p* = 0.005, η^2^□ = 0.03, Figure 5A), alpha (F_3,263_ = 4.98 to 5.56, *p* = 0.001 to 0.0007, η^2^□ = 0.02 to 0.03, Figure 5B), and beta (F_3,263_ = 3.80, *p* = 0.042, η^2^□ = 0.02, Figure 5C) bands. For theta and alpha bands, the effects were driven by lower values during the visualization of the knee-position than real-body visualization (*p* = 0.033 to 0.003, *d* = 0.32 to 0.43), while no-body visualization was intermediate. Regarding beta band, the effect was only due to a tendency of lower pre-event values during the visualization of knee-position cue than no-body visualization (*p* = 0.06, *d* = 0.31). We further identified post-event body visualization effect in theta (F_3,263_ = 3.61, *p* = 0.02, η^2^□ = 0.02), alpha (F_3,263_ = 3.36 p = 0.03, η^2^□ = 0.03), and beta (F_3,263_ = 3.47, *p* = 0.042, η^2^□ = 0.02) bands. For theta band, the effects were again driven by lower values during the visualization of the knee-position cue compared to real-body visualization (*p* = 0.033, *d* = 0.31), while no-body visualization was intermediate, and nothing subsisted for beta band. As for alpha an interaction was identified (F_6,263_ = 2.23, *p* = 0.043, η^2^□ = 0.04), driven by the motor-enhanced level, where the visualization of knee-position cue showed lower values than real-body visualization (*p* = 0.01, *d* = 0.39). All body visualization effects were not robust to sensitivity analyses (F_2,263_ = 0.35 to 2.71, *p* = 1). Baseline differences (F_2,263_ = 2.87 to 7.71, *p* = 0.04 to 0.0007, η^2^□ = 0.02 to 0.17) was driven by participants exhibiting more power during the visualization of knee-position cue (theta, alpha and beta bands) than real-body and no-body visualization (supplementary Figure S1).

**Figure 5.**
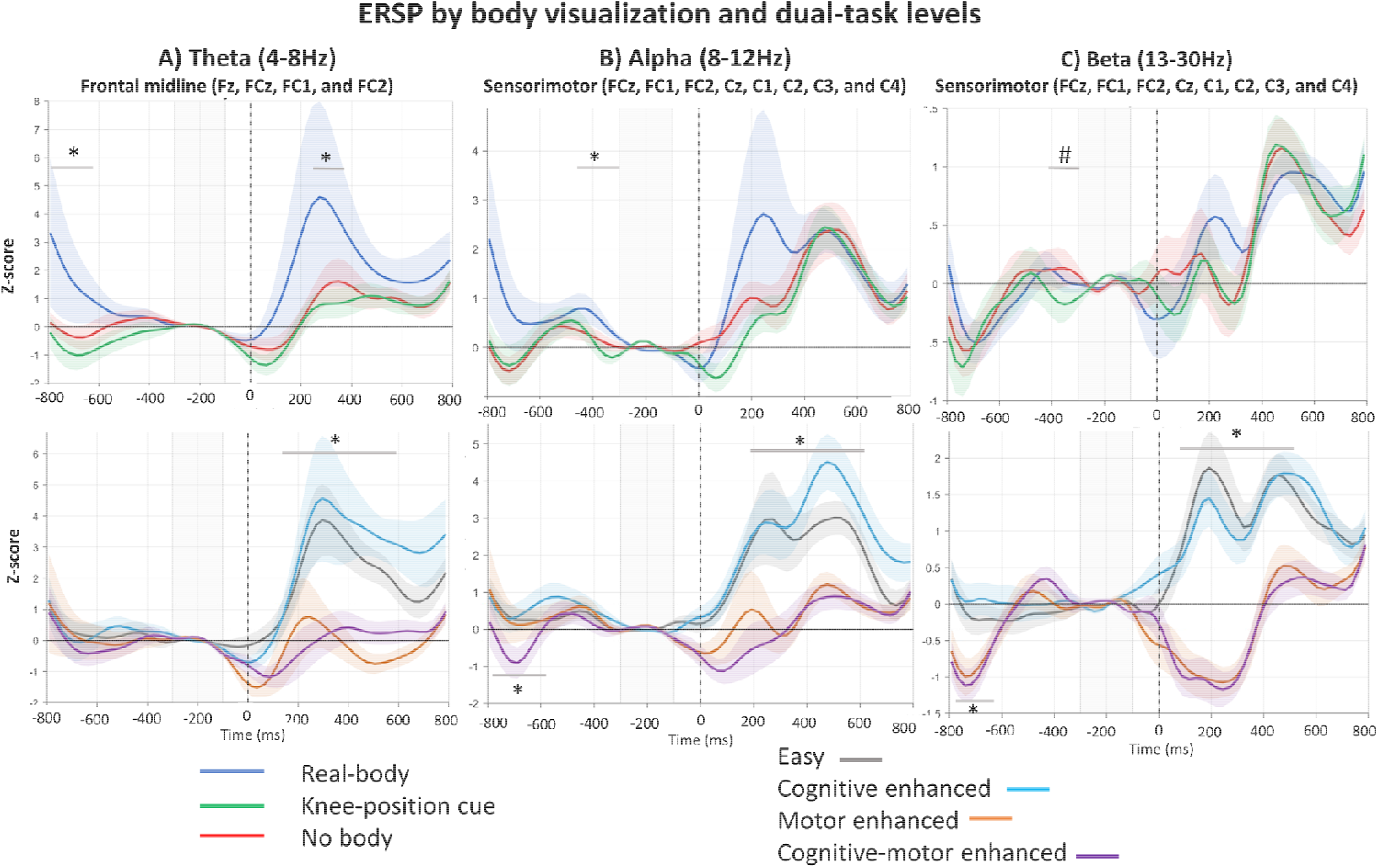
Event-related spectral perturbation (ERSP) across body visualization conditions and dual-task levels in (A) frontal midline theta, (B) sensorimotor alpha, and (C) sensorimotor beta bands. Panels show baseline-corrected spectral power expressed as Z-scores relative to the −300 to −100 ms baseline period, with corresponding standard errors of the mean (SEM). **Time zero represents the moment of clearing the obstacles during the stepping execution**. Asterisks indicate time windows with significant (*p* < 0.05) pairwise differences identified by cluster-based permutation testing, whereas hashtags indicate statistical trends (*p* < 0.09). For body visualization, **(A)** theta power was lower, during the visualization of knee-position than real-body visualization both before and after stepping, with the no-body visualization condition showing an intermediate response. **(B)** In the alpha band, the visualization of knee-position cue similarly showed lower power than real-body visualization, but only after the stepping. **(C)** In the beta band, only a trend-level difference between no-body visualization and that of knee-position cue was observed before stepping. For dual-task level, post-event effects were observed across all three frequency bands **(A–C):** the easy and cognitive-enhanced levels showed comparable power, both differing from the motor-enhanced and cognitive–motor-enhanced levels. Pre-event effects were observed in the alpha and beta bands (B–C), with lower power in the motor-enhanced and cognitive–motor-enhanced levels than in the easy and cognitive-enhanced levels. Only the effects of dual-task level remained robust in sensitivity analyses of raw power when baseline activity was included as a covariate.

We identified a post-event main effect of dual-task levels in all frequency bands (F_3,263_ = 3.20 to 22.00, *p* = 0.037 to 0.0001, η^2^□ = 0.02 to 0.17, Figure 5). This manifested through an increase in the easy and cognitive-enhanced conditions on the one hand compared to the motor-enhanced and cognitive-motor-enhanced levels on the other hand (*p* = 0.018 to 0.001; *d* = 0.34 to 0.72). A similar pattern was observed for pre-event period with this time lower values in the motor-enhanced and cognitive-motor-enhanced levels (*p* = 0.047 to 0.006; *d* = 0.32 to 0.48) but only for alpha and beta bands. After sensitivity analyses, the dual-task level effects in all bands subsisted (F_3,263_ = 10.98 to 37.71, *p* = 0.001 to 0.0001, η^2^□ = 0.03 to 0.29) despite baseline difference only in theta band as motor-enhanced and cognitive-motor-enhanced exhibited higher baseline power than easy and cognitive-enhanced levels (supplementary Figure S1).

## 4 Discussion

The present study investigated the impact of different scenarios of body visualization in XR on cognitive-motor adaptation during an obstacle avoidance stepping task. The main finding was that while the absence of body visualization degraded motor performance, adding a cue on knee position restored it, by increasing stepping accuracy and reducing omissions, to levels comparable to when participants visualized their real body. Interestingly, at the neural level, markers of conflict monitoring (N450), sensory integration and motor preparation (sensorimotor alpha and beta activities), co-occurred with these benefits of cueing the knee position. Moreover, we did not detect improvement in embodiment or sense of presence accompanying the behavioral benefits of cueing compared with the absence of body visualization. Finally, cognitive performance was primarily affected by the attentional demands of the task rather than body visualization.

### 4.1 A cue on the knee position compensates for the degradation of stepping performance induced by the absence of body visualization

In line with our principal hypothesis, participants exhibited similar stepping accuracy and omission during real-body visualization and that of the knee-position cue, with both conditions yielding better outcomes than no-body visualization. These observations support the degradation of movement quality in the absence of body visualization compared to partial or full-body visualization as previously reported (20, 21).

The present study contributes new knowledge to the existing literature by showing that a minimal body representation, limited to a visual cue on the knee position, is sufficient to restore motor performance to the level comparable to that achieved with direct visualization of the real body. Previous reports argued that body visualization would impact performance through the availability, precision, and congruence of body-related sensory feedback (13–16). Our interpretation is that the cue on knee position likely increased the precision of body-related information by providing participants with a reliable estimate of the distance needed to step over the obstacle which was not possible in the absence body visualization. This assumption is grounded in previous work where visual feedback on lower extremities improves performance in similar obstacle avoidance tasks (64, 65).

Interestingly, the benefit of the knee-position cue occurred despite the absence of any increase in sense of embodiment or presence. Indeed, in the absence of body visualization or during the visualization of knee-position cue, participants reported less sense of embodiment and presence compared with real-body visualization. However, there was no difference in participants’ self-reported experiences between no-body visualization and that of the knee-position cue. These observations suggest that the benefits of cueing in its current form could operate independently of participants’ self-reported experience, as previously shown (14). It introduces the idea that task-relevant body-related feedback plays the most critical role in guiding the movement. Indeed, when body visualization is manipulated in XR, the resulting disruption of sensorimotor information appears to be the primary driver of its effects (14), which may not necessarily be mediated by changes in the sense of embodiment (13, 15, 16). Moreover, the stronger embodiment during the direct perception of one’s real body than when the body is only partially represented or completely absent is consistent with works on bodily embodiment. Indeed, a richer and more coherent body representation, aligned with the available sensory information, would enhance the components of embodiment (12, 66, 67).

The benefit of cueing was associated with enhanced conflict monitoring particularly in the high dual-task levels. Conflict monitoring is the ability to detect and/or resolve interference arising from competing response tendencies or task-relevant information (68). The N450 component in ERP is a well-established neural marker of conflict monitoring in tasks with important interference (43, 44). Our study showed an interaction effect exhibiting a smaller N450 amplitude during no-body than real-body visualization, whereas the visualization of knee-position cue showed an intermediate pattern. This result is consistent with our hypothesis and is in favor of a potential facilitation of conflict monitoring when richer and more congruent feedback is present in real-body visualization compared to no-body visualization. Moreover, we interpret the intermediate N450 response during the visualization of knee-position cue as a potential restoration of the capacity of conflict monitoring. Indeed, it is possible that the information provided by the knee-position cue on the distance required to clear the obstacle might have reduced the competition between the two tasks, which is not possible during no-body visualization. Moreover, as indicated by interaction effect, body visualization impacted the marker of conflict monitoring particularly at the highest dual-task levels, where conflict is likely to be the most important. However, the fact that there was no interaction effect on behavioral performance regarding the benefit of cueing can be intriguing. Nevertheless, these two phenomena are not necessarily contradictory. It might be that the potential restoration of conflict monitoring capacity occurring during the high dual-task level would reflect a strategy that supports stable performance across levels. This could be associated with a dynamic reallocation of resources to motor execution, consistent with the capacity-sharing model of dual-task paradigms (29).

The preparation and selection of motor strategies may constitute other mechanisms through which body visualization influences behavioral performance. Indeed, the visual feedback available to guide movement can shape motor planning and the selection of appropriate strategies (14, 16). Changes in sensorimotor alpha-and beta-band activity have been widely used to investigate these processes, including during dual-task performance (40, 46, 61, 69). ERSPs in the present study revealed interesting observations: the visualization of the knee-position cue elicited lower alpha values than real-body visualization, whereas no-body and real-body visualizations did not differ. We interpret this as the role of alpha event-related desynchronization (ERD) before movement which is associated with the increase in cortical excitability (60, 62) or the capacity of sensory integration (69). This pattern on alpha has been identified even especially in similar obstacle avoidance tasks (70, 71). It is thus possible that to benefit from the knee-position cue, participants needed an increase in cortical excitability and/or sensory integration. This is supported by research showing that a visual cue elicited strong alpha ERD in the sensorimotor areas during motor preparation (72). Moreover, post-event effects in alpha band (*i.e*., lower values during real-body visualization than that of knee-position cue), might indicate prolonged excitability and sensorimotor integration after movement (73). Indeed, a diminution of alpha power post-movement can be a result of a less complete return to the idling state to anticipate the next movement (73). In this paradigm, it is consistent with the benefits of cueing, as the reliance on visual cues to continuously update body-related information may require anticipatory processing given the continuous nature of the task. The limited effects observed in the beta band may appear surprising, given that our primary hypothesis on motor strategies concerned beta-band modulation. However, we interpret this finding reflecting that body visualization does not directly influence the core processes of motor preparation, consistent with the established role of beta-band activity in the release of motor inhibition. Instead, body visualization may primarily affect the integration of sensory information available to the motor system, rather than the motor preparatory mechanisms themselves. This assumption is further supported by the fact that the effects of body visualization have been associated with the congruence, availability and precision of sensory feedback (13-16) which is coherent with the role of alpha band modulation (57,59, 67).

As our paradigm involves dual-tasking, it requires an adaptation of cognitive control to allocate attentional resources. Thereby, frontal midline theta activity is a well-established neural marker of cognitive control and has been extensively used to assess attentional demands (58, 74). Contrary to our expectations, the effect of cueing was also reflected in the theta band, with lower values both pre-and-post-event during the visualization of the knee-position cue than real-body visualization. This could be interpreted as theta ERD suggesting that the presence of the cue reduced overall cognitive control during movement. At first glance a theta ERD during a dual-task appears counterintuitive, as frontal midline theta activity is typically reported to increase during cognitive control and especially during conflict monitoring (58). An explanation may arise in the results of sensitivity analyses indicating that the effects of body visualization on oscillation dynamics depend on baseline activity. Indeed, frontal midline theta is known to reach its maximum during periods of high conflict (58). In our paradigm, conflict may have peaked immediately before stepping initiation, suggesting that theta activity was already elevated during the baseline period. Consequently, the apparent ERD, rather than the more commonly reported event-related synchronization, may simply reflect a high baseline theta level rather than a true event-related reduction in theta activity. This interpretation aligns with the real-body visualization exhibiting more baseline theta power than no body visualization and the visualization of knee-position cue being intermediary (supplementary Figure S1). This might suggest that the benefits of cueing, at least in its current implementation, are not obtained without a cost. Indeed, they do not rely solely on pure conflict monitoring and motor preparation as we hypothesized. There might be an increase in the overall need for cognitive control as suggested by theta dynamics reported here.

### 4.2 The dual-task level impacted both cognitive and stepping performance while body visualization impacted only motor changes

The improvement in stepping performance with increasing dual-task levels may appear counterintuitive as we hypothesized the inverse. One possible explanation is the fixed order of the levels which may have induced a learning effect within a session. Interestingly, because previous studies used a fixed task order yet still reported CMI on performance (30), these learning effects are unlikely to be solely attributable to the fixed order. Indeed, our sample consisted of highly physically active young adults with well-developed motor abilities. Therefore, the stepping task might have been relatively simple, making it likely to become at least partially automatic for this population. Under these conditions, it is possible that the effects of CMI on motor performance were minimized. Indeed, it is known that to preserve their performance during dual-task, young adults can adopt motor automation strategies when the motor task is sufficiently simple (75). This interpretation further aligns with the absence of an interaction effect between body visualization and dual-task level and the absence of an effect on the dual-task cost on stepping performance. It also suggests that the benefit of cueing remained stable across dual-task levels because the stepping task did not become sufficiently demanding.

In contrast, cognitive performance declined as dual-task level increased as anticipated, but regardless of the body visualization. Interestingly, there was not an effect of body visualization on cognitive performance. It is plausible that body-related feedback mainly impacted motor execution rather than cognition, through the modulation of sensorimotor feedback, consistent with previous studies in virtual reality (76), where increasing attentional demands preferentially modulated cognitive performance. Therefore, participants in the present study were able to preserve motor performance, whether aided by the knee-position cue or not, at the expense of cognitive performance. This reveals an interesting dissociation in our paradigm: body visualization primarily influenced stepping performance, whereas the level of CMI predominantly affected cognitive performance. Previous findings showed that young adults can preserve both motor and cognitive performances while relying on compensatory mechanisms during dual-tasks (29). In the present study, fact that motor performance was facilitated while cognitive performance declined may reflect that during CMI the motor task was sufficiently simple to prioritize and maintain (77). It is conceivable that increasing the motor load, by further increasing the task’s complexity for example, would exceed the available compensatory capacity therefore allowing to investigate further dual-task related modulation performances.

Interestingly, increasing dual-task levels was also associated with modulations of conflict monitoring and resource allocation markers. Specifically, higher dual-task levels elicited larger N450 and smaller P300 amplitudes. This pattern is consistent with markers of increased CMI, whereby greater task demands increase conflict monitoring while limiting the capacity to share attentional resources between the two tasks (43, 45, 46, 55). Therefore, the dual-task manipulation, particularly the cognitive load, successfully induced CMI and neural resources were shared between cognitive and movement execution. The results also complement our previous assumption that the motor task was easy for this population. Indeed, young adults could have preserved motor performance also because of compensatory mechanisms allowing a priority allocation of attentional resources toward movement execution (29). Thereby, both the easiness of the task and compensatory mechanisms reflected in the concurrent increase in N450 and decrease in P300 could have supported the facilitation observed for stepping performance. The potential redistribution of resources was also reflected in the oscillatory activities. Increasing dual-task levels was associated with a decrease in pre-event alpha and beta power, indicating greater engagement of sensorimotor networks during movement preparation. In parallel, the post-movement value was reduced at higher dual-task levels, suggesting that the sensorimotor cortex returned less completely to its post-movement idling state in anticipation of the next stepping execution (78).

The effects of the dual-task levels on overall cognitive control, as displayed by theta activity, deserve particular attention and complement our previous interpretation regarding the effects of body visualization. Like the alpha and beta bands, post-movement theta values were reduced at higher levels of dual-task, contrary to the well-known increase. In our paradigm, theta activity was already elevated during the pre-event baseline at higher dual-task levels (supplementary Figure S1), indicating that participants entered movement preparation with a high level of cognitive engagement. Previous work shown that frontal midline theta does not necessarily increase linearly with cognitive demands and may approach a functional ceiling under conditions of sustained conflict (74). Thus, one possible interpretation is that participants operating under the highest dual-task levels had already recruited cognitive control processes to a near-maximal level before movement onset, leaving limited capacity for an additional theta increase.

### 4.3 Implication for XR community and future work

Taken together, the effects of body visualization on motor performance support the incorporation of body-related visual cues in XR when preserving motor performance is a primary objective. Depending on the context, it may be sufficient to provide task-relevant visual body cues rather than a complete body representation. This should inform XR design as more sophisticated solutions, such as partial or full avatars, and the real body visualization can be time and resource consuming. If behavioral performance is the primary objective, our findings suggest that task-relevant body cues may be sufficient, thus simplifying the design process. Moreover, our findings emphasize that the XR features could differently impact behavioral performance and the self-reported user experience. In case of body visualization, the information required to efficiently guide movement, and performance might not necessarily be the same as that required to enhance embodiment, presence, and overall user experience. Consequently, the choice of body representation should be driven by the intended objectives (23, 25) rather than by the sole assumption that richer representations are inherently superior regardless of the end goal.

Neural markers offer complementary insights into the processes underlying the benefits of the knee-position cue, suggesting the involvement of motor preparation, sensorimotor integration, and enhanced conflict monitoring. Thereby, it opens several promising avenues for future research. Beyond simple visual cues, one should investigate how progressively enriching body feedback, through avatars, varying levels of visual realism, or manipulating the meaning and affordances of virtual objects with respect to the body, influences behavior and its underlying processes. Such manipulations would not only inform the design of more effective XR environments but also provide valuable insights into the fundamental role of body-related feedback in perception and action.

Furthermore, the results are particularly relevant for clinical practice in rehabilitation, where fully immersive virtual reality may be preferable to mixed reality in certain contexts because it provides complete control over the visual environment and therapeutic contents (79). For example, in the assessment and rehabilitation of spatial neglect following stroke, considerable efforts are currently being devoted to leveraging XR technologies to manipulate visual and spatial information in ways that are not feasible in the real world (80). Building on our findings, incorporating simple body-related visual cues could preserve motor performance while retaining the flexibility and experimental control afforded by fully immersive virtual environments.

### 4.4 Limitations

The main limitation of this study is that the stepping task was relatively simple for our sample of young, physically active participants, which may have limited our ability to fully investigate the effects of body visualization under more demanding dual-task conditions. Nevertheless, even within this relatively easy task, manipulating body visualization, through a minimal cue on knee position, produced measurable changes in both motor performance and its underlying cerebral bases. It highlights the importance of body-related sensory information in XR and suggests that its integration should be carefully considered whenever preserving motor performance is a key objective. In addition, given the sample’s characteristics, the generalizability of the results to older adults or clinical populations remains to be established in future studies.

The results regarding the motor preparation or sensory integration may reflect a more tonic modulation of neural activity rather than a phasic event-related response since none of the effects of body visualization in ERSPs survived the sensitivity analyses. This apparent tonic modulation may stem from several characteristics of the experimental paradigm. The task was continuous, with relatively short inter-trial intervals, potentially limiting the return to baseline and the emergence of distinct event-related oscillatory responses. Furthermore, trial-by-trial feedback on performance may have maintained sustained cognitive engagement and monitoring throughout the task. To further complement the information on oscillatory dynamics underlying these effects, we performed an exploratory analysis on raw time frequency power. Interestingly, raw power was modulated throughout the trial, with differences between conditions showing the same directionality both before and after the event, as well as during the baseline period (supplementary Figure S1). This pattern supports the assumption of the tonic effects and further mirrors ERSP observations, since the visualization of knee-position cue exhibited changes in alpha and beta activity consistent with increased cortical excitability, sensory integration, and motor preparation (supplementary Figure S1). Future studies with dedicated event construction would be useful to delve deeper into these processes.

One could therefore question why we did not select a baseline period that would be more stable and better suited to isolate event-related activity. However, identifying such a stable period in the current version of our task is particularly challenging. Because the task is continuous, with occasionally short inter-trial intervals and overlapping cognitive demands, periods that could be considered free from task-related processing are difficult to define. These characteristics likely contributed to the tonic modulation observed in the present data. We chose the pre-event period as our reference, as this time window is consistently described in the literature for ERSPs (40). While this choice does not fully disentangle tonic from phasic activity, it provides a physiologically meaningful reference that is directly comparable with previous studies. With these initial findings, we are now in a better position to refine the task design. Future iterations will include modifications to the task structure to create more suitable baseline periods and more appropriate event detection, while also increasing the complexity of the motor task.

Finally, technologies used to provide XR environments do not have the same technical characteristics. The projection of the real environment onto a screen through the headset’s embedded cameras may lead to variations in image resolution, refresh rate, field of view, and most importantly image distortion. Thus, the impact on the fidelity and congruence of body-related feedback may vary. The present results might therefore be limited to the Meta Quest 3 technology which could be attenuated with a lower-specification device or amplified with a higher-specification one. Nevertheless, it has been reported that Meta Quest 3 has an important added value in both rehabilitation and basic science (81, 82), strengthening the direct application of our findings.

## 5 Conclusion

This original research investigated cognitive–motor adaptation in XR while manipulating body visualization and simultaneously assessing its cerebral bases using mobile brain/body imaging. Both behavioral and neural changes support the benefit of task-relevant cues to preserve cognitive-motor performance. Importantly, the results suggest that effective body representation in XR does not necessarily require realistic or complete visualization; rather, minimal cues may provide sufficient information to support cognitive-motor adaptation. This finding has direct implications for XR design, emphasizing that the nature and amount of body-related information presented to users should be tailored to the intended objectives. Beyond these practical implications, our results open promising avenues for investigating how specific features of immersive environments shape cognitive-motor behavior and its underlying mechanisms. The integration of XR and MoBI may provide a powerful framework for understanding how the users adapt to immersive environments while leveraging well-known neural markers of cognitive-motor control. Ultimately, this approach could help move XR toward more evidence-based and personalized usages, in which immersive features are designed according to users’ sensorimotor and cognitive requirements to maximize their benefits in research, healthcare, and rehabilitation.

## 7 Data availability statement

Behavioral data, EEG data, and code used for analyses are publicly available in the following repository. https://osf.io/dy58u/overview?view_only=52b672d1b1c54c3f9cf6064a6f625f58

## 8 Acknowledgments

The authors thank Alexandre Nahon, Brayan Silliau, and Cédric Sockeng Kofo for their involvement in the design of the cognitive-motor stepping task. The authors also thank all the participants who took part in the investigations.

