## Supplementary File for "Uncovering cognitive-motor adaptation to body visualization in immersive technologies through their behavioral and electrophysiological correlates"

^2^3iL Ingénieurs, Limoges, France

^3^Université Côte d′Azur, LAMHESS, Nice, France

* Corresponding author: Elpidio Attoh-Mensah

Address: 123 Avenue Albert Thomas, 87000, Limoges

Table S1. Behavioral performances

|  | Easy Stepping_ST | 1-Back _ST | Easy DT | 2-back_ST | Cognitive-enhanced_DT | Hard Stepping_ST | Motor-enhanced_DT | Cognitive-motor_enhanced_DT |
| --- | --- | --- | --- | --- | --- | --- | --- | --- |
| Stepping accuracy (%) | | | | | | | | |
| Real Body | 86.9 (16.7) | NA | 87.9 (14.0) | NA | 85.5 (17.14) | 90.7 (11.8) | 94.5 (12.2) | 95.4 (9.9) |
| Knee-position cue | 89.8 (13.7) | NA | 87.2 (15.9) | NA | 89.0 (15.7) | 89.5 (20.5) | 91.8 (12.2) | 94.2 (9.3) |
| No Body | 81.5 (19.3) | NA | 81.8 (18.7) | NA | 87.3 (16.4) | 88.6 (14.9) | 89.4 (14.9) | 89.9 (15.2) |
| Stepping omission | | | | | | | | |
| Real Body | 8.5 (12.2) | NA | 7.8 (10.2) | NA | 9.5 (12.5) | 11.5 (17.2) | 6.5 (17.10) | 5.3 (14.8) |
| Knee-position cue | 6.5 (10.2) | NA | 8.4 (11.5) | NA | 7.10 (11.4) | 9.48 (16.3) | 9.7 (16.3) | 6.6 (12.7) |
| No Body | 12.6 (14.2) | NA | 12.4 (13.5) | NA | 8.3 (11.9) | 15.10 (21.7) | 14.0 (21.4) | 13.3 (22.0) |
| N-back accuracy (%) | | | | | | | | |
| Real Body | NA | 96.8 (5.6) | 90.3 (12.5) | 75.6 (21.3) | 71.4 (20.5) | NA | 87.3 (17.4) | 72.1 (21.6) |
| Knee-position cue | NA | 96.4 (8.9) | 90.7 (13.1) | 77.0 (20.14) | 72.1 (22.7) | NA | 88.07 (13.5) | 69.7 (23.3) |
| No Body | NA | 94.6 (9.9) | 89.6 (12.4) | 74.7 (20.7) | 69.0 (21.5) | NA | 87.8 (14.6) | 69.6 (19.7) |
| N-back omission | | | | | | | | |
| Real Body | NA | 3.5 (13.8) | 5.9 (14.9) | 10.8 (18.6) | 12.8 (19.4) | NA | 6.3 (16.7) | 12.0 (19.8) |
| Knee-position cue | NA | 3.2 (13.6) | 5.9 (15.5) | 10.5 (18.5) | 11.9 (20.6) | NA | 6.3 (15.8) | 12.2 (20.4) |
| No Body | NA | 3.7 (14.7) | 6.2 (14.7) | 11.2 (18.4) | 13.2 (19.8) | NA | 6.3 (15.6) | 13.4 (18.6) |

Values are mean with corresponding standard deviation in parentheses

Table S2. Exact windows and length for the main effects for ERP

| ROI | Effect | Families | Start_ms | End_ms | Duration_ms | Mean_Fval | Mean_pval | Min_pval | Mean_partal_eta |
| --- | --- | --- | --- | --- | --- | --- | --- | --- | --- |
| Fronto-Central | Main dual-task level | Dual-tasl level | 292 | 500 | 208 | 6,957108929 | 0,00019996 | 0,00019996 | 0,064824866 |
| Fronto-Central | Interaction | Interaction | 376 | 500 | 124 | 2,420388661 | 0,031793641 | 0,031793641 | 0,046164999 |
| Centro-Parietal | Main dual-task level | Dual-tasl level | 416 | 600 | 184 | 5,066608607 | 0,00239952 | 0,00239952 | 0,048025082 |

| ROI | Effect | Families | Start_ms | End_ms | Duration_ms | Mean_Fval | Mean_pval | Min_pval | Mean_partal_eta | Window |
| --- | --- | --- | --- | --- | --- | --- | --- | --- | --- | --- |
| Frontocentral | Main body visualization | Body visualization | -792 | -584 | 208 | 5,640754428 | 0,007109689 | 0,00239952 | 0,036199682 | PreMovement |
| Frontocentral | Main body visualization | Body visualization | 88 | 372 | 284 | 3,653272129 | 0,020145971 | 0,0059988 | 0,023763241 | PostMovement |
| Frontocentral | Main dual-task level | Dual-task level | 12 | 36 | 24 | 3,010328161 | 0,036992601 | 0,035392921 | 0,029223278 | PostMovement |
| Frontocentral | Main dual-task level | Dual-task level | 244 | 788 | 544 | 6,838375974 | 0,001681482 | 0,00019996 | 0,063054125 | PostMovement |
| Sensorimotor | Main body visualization | Body visualization | -792 | -688 | 104 | 4,638846231 | 0,014237153 | 0,0019996 | 0,029975879 | PreMovement |
| Sensorimotor | Main body visualization | Body visualization | -376 | -328 | 48 | 5,568039363 | 0,011531027 | 0,00139972 | 0,035730969 | PreMovement |
| Sensorimotor | Main dual-task level | Dual-task level | -688 | -688 | 0 | 2,926265484 | 0,049990002 | 0,049990002 | 0,028430697 | PreMovement |
| Sensorimotor | Main dual-task level | Dual-task level | -92 | -16 | 76 | 4,374924457 | 0,00959808 | 0,00619876 | 0,041909228 | PreMovement |
| Sensorimotor | Main body visualization | Body visualization | 116 | 140 | 24 | 3,009801951 | 0,048490302 | 0,046990602 | 0,019670625 | PostMovement |
| Sensorimotor | Main body visualization | Body visualization | 192 | 296 | 104 | 3,581834817 | 0,025794841 | 0,016396721 | 0,023319927 | PostMovement |
| Sensorimotor | Main dual-task level | Dual-task level | 12 | 764 | 752 | 10,85725196 | 0,001846297 | 0,00019996 | 0,0952305 | PostMovement |
| Sensorimotor | Interaction | Interaction | 88 | 116 | 28 | 2,238373085 | 0,035892821 | 0,035392921 | 0,042848085 | PostMovement |
| Sensorimotor | Main body visualization | Body visualization | -792 | -792 | 0 | 3,333402941 | 0,046790642 | 0,046790642 | 0,021739575 | PreMovement |
| Sensorimotor | Main body visualization | Body visualization | -328 | -328 | 0 | 3,808913091 | 0,033793241 | 0,033793241 | 0,02476393 | PreMovement |
| Sensorimotor | Main dual-task level | Dual-task level | -792 | -636 | 156 | 8,443804349 | 0,006027366 | 0,00019996 | 0,07725391 | PreMovement |
| Sensorimotor | Main dual-task level | Dual-task level | -92 | -16 | 76 | 4,757868324 | 0,00844831 | 0,00079984 | 0,045337335 | PreMovement |
| Sensorimotor | Main body visualization | Body visualization | 272 | 272 | 0 | 3,47425776 | 0,046190762 | 0,046190762 | 0,022637397 | PostMovement |
| Sensorimotor | Main dual-task level | Dual-task level | 12 | 712 | 700 | 22,00667344 | 0,001706801 | 0,00019996 | 0,170654506 | PostMovement |

Table S3. Exact windows and length for the main effects for ERSPs

Table S4. Exact windows and length for pairwise comparison for ERP

| ROI | Contrast | Families | Start_ms | End_ms | Duration_ms | Mean_tval | Mean_pval | Min_pval | Cohen_d |
| --- | --- | --- | --- | --- | --- | --- | --- | --- | --- |
| Fronto-Central | Easy_DT vs Motor_Enhanced_DT | Bloc | 292 | 472 | 180 | 3,808944961 | 0,00504247 | 0,00019996 | 0,43981908 |
| Fronto-Central | Easy_DT vs Cognitive-Motor_Enhanced_DT | Bloc | 320 | 416 | 96 | 3,170938597 | 0,022579484 | 0,00619876 | 0,36614845 |
| Fronto-Central | Cognitively_Enhanced_DT vs Motor_Enhanced_DT | Bloc | 296 | 480 | 184 | 3,613207264 | 0,006096653 | 0,00019996 | 0,417217237 |
| Fronto-Central | Cognitively_Enhanced_DT vs Cognitive-Motor_Enhanced_DT | Bloc | 336 | 388 | 52 | 3,005239878 | 0,028251493 | 0,01219756 | 0,347015211 |
| Fronto-Central | Cognitively_Enhanced_DT vs Cognitive-Motor_Enhanced_DT | Bloc | 460 | 468 | 8 | 2,842042446 | 0,042658135 | 0,033993201 | 0,328170794 |
| Centro-Parietal | Easy_DT vs Motor_Enhanced_DT | Bloc | 480 | 508 | 28 | 2,984797359 | 0,022045591 | 0,01219756 | 0,344654712 |
| Centro-Parietal | Easy_DT vs Cognitive-Motor_Enhanced_DT | Bloc | 460 | 464 | 4 | 2,620330453 | 0,047890422 | 0,046190762 | 0,302569698 |
| Centro-Parietal | Easy_DT vs Cognitive-Motor_Enhanced_DT | Bloc | 472 | 600 | 128 | 2,793322982 | 0,032466234 | 0,01179764 | 0,322545155 |
| Centro-Parietal | Cognitively_Enhanced_DT vs Motor_Enhanced_DT | Bloc | 464 | 532 | 68 | 3,298913267 | 0,016963274 | 0,00079984 | 0,380925693 |
| Centro-Parietal | Cognitively_Enhanced_DT vs Cognitive-Motor_Enhanced_DT | Bloc | 456 | 600 | 144 | 3,384403074 | 0,007587672 | 0,00119976 | 0,390797205 |
| Fronto-Central | REAL BODY vs NO BODY @ Motor_Enhanced_DT | Interaction | 468 | 468 | 0 | -2,941407442 | 0,045590882 | 0,045590882 | -0,339644476 |
| Fronto-Central | REAL BODY vs NO BODY @ Cognitive-Motor_Enhanced_DT | Interaction | 496 | 496 | 0 | -2,918747425 | 0,049590082 | 0,049590082 | -0,337027922 |
| Fronto-Central | REAL BODY vs NO BODY @ Motor_Enhanced_DT | Interaction | 444 | 444 | 0 | -2,711182833 | 0,084383123 | 0,084383123 | -0,313060428 |
| Fronto-Central | REAL BODY vs NO BODY @ Motor_Enhanced_DT | Interaction | 460 | 460 | 0 | -2,780282259 | 0,069786043 | 0,069786043 | -0,321039342 |
| Fronto-Central | REAL BODY vs NO BODY @ Cognitive-Motor_Enhanced_DT | Interaction | 492 | 492 | 0 | -2,749558687 | 0,079184163 | 0,079184163 | -0,31749169 |
| Fronto-Central | REAL BODY vs NO BODY @ Cognitive-Motor_Enhanced_DT | Interaction | 500 | 500 | 0 | -2,762582064 | 0,077184563 | 0,077184563 | -0,3189955 |

Table S5. Exact windows and length for pairwise comparison for ERSPs

| ROI | Contrast | Families | Start_ms | End_ms | Duration_ms | Mean_tval | Mean_pval | Cohen_d | Band |
| --- | --- | --- | --- | --- | --- | --- | --- | --- | --- |
| Frontocentral | REAL BODY vs KNEE-POSITION CUE | Body visualization | -792 | -612 | 180 | 3,26986176 | 0,017271546 | 0,377571113 | theta |
| Frontocentral | Easy_DT vs Motor_Enhanced_DT | Dual-task Level | -16 | -16 | 0 | 2,89456169 | 0,032593481 | 0,334235194 | theta |
| Frontocentral | REAL BODY vs KNEE-POSITION CUE | Body visualization | 220 | 296 | 76 | 2,673045674 | 0,038342332 | 0,308656728 | theta |
| Frontocentral | Easy_DT vs Motor_Enhanced_DT | Dual-task Level | 12 | 36 | 24 | 2,941229127 | 0,015396921 | 0,339623886 | theta |
| Frontocentral | Easy_DT vs Motor_Enhanced_DT | Dual-task Level | 372 | 608 | 236 | 3,544016992 | 0,009018196 | 0,409227833 | theta |
| Frontocentral | Easy_DT vs Cognitive-Motor_Enhanced_DT | Dual-task Level | 452 | 528 | 76 | 2,840794791 | 0,025744851 | 0,328026727 | theta |
| Frontocentral | Cognitively_Enhanced_DT vs Motor_Enhanced_DT | Dual-task Level | 324 | 788 | 464 | 4,021067937 | 0,007587956 | 0,464312931 | theta |
| Frontocentral | Cognitively_Enhanced_DT vs Cognitive-Motor_Enhanced_DT | Dual-task Level | 272 | 788 | 516 | 3,27261753 | 0,016710944 | 0,377889322 | theta |
| Sensorimotor | REAL BODY vs NO BODY | Body visualization | -792 | -740 | 52 | 2,866618961 | 0,045190962 | 0,331008646 | alpha |
| Sensorimotor | REAL BODY vs KNEE-POSITION CUE | Body visualization | -352 | -328 | 24 | 3,541900253 | 0,00739852 | 0,408983413 | alpha |
| Sensorimotor | Easy_DT vs Motor_Enhanced_DT | Dual-task Level | -92 | -68 | 24 | 2,988820144 | 0,034493101 | 0,345119223 | alpha |
| Sensorimotor | Cognitively_Enhanced_DT vs Cognitive-Motor_Enhanced_DT | Dual-task Level | -16 | -16 | 0 | 2,959895808 | 0,042791442 | 0,341779328 | alpha |
| Sensorimotor | Easy_DT vs Cognitively_Enhanced_DT | Dual-task Level | 452 | 504 | 52 | -2,9716563 | 0,035259615 | -0,343137313 | alpha |
| Sensorimotor | Easy_DT vs Motor_Enhanced_DT | Dual-task Level | 244 | 608 | 364 | 3,675703001 | 0,005958808 | 0,424433623 | alpha |
| Sensorimotor | Easy_DT vs Cognitive-Motor_Enhanced_DT | Dual-task Level | 12 | 608 | 596 | 3,657732927 | 0,007656802 | 0,422358618 | alpha |
| Sensorimotor | Cognitively_Enhanced_DT vs Motor_Enhanced_DT | Dual-task Level | 12 | 12 | 0 | 2,86441348 | 0,048590282 | 0,330753979 | alpha |
| Sensorimotor | Cognitively_Enhanced_DT vs Motor_Enhanced_DT | Dual-task Level | 272 | 736 | 464 | 5,017965782 | 0,003683474 | 0,579424779 | alpha |
| Sensorimotor | Cognitively_Enhanced_DT vs Cognitive-Motor_Enhanced_DT | Dual-task Level | 12 | 736 | 724 | 4,60614818 | 0,003723393 | 0,531872178 | alpha |
| Sensorimotor | REAL BODY vs KNEE-POSITION CUE @ Motor_Enhanced_DT | Interaction | 64 | 192 | 128 | 3,442569141 | 0,017996401 | 0,397513644 | alpha |
| Sensorimotor | NO BODY vs KNEE-POSITION CUE | Condition | -328 | -328 | 0 | 2,746291361 | 0,075784843 | 0,317114411 | beta |
| Sensorimotor | Easy_DT vs Motor_Enhanced_DT | Dual-task Level | -792 | -716 | 76 | 3,451657281 | 0,01174765 | 0,398563052 | beta |
| Sensorimotor | Easy_DT vs Cognitive-Motor_Enhanced_DT | Dual-task Level | -792 | -688 | 104 | 3,784907762 | 0,008878224 | 0,437043503 | beta |
| Sensorimotor | Cognitively_Enhanced_DT vs Motor_Enhanced_DT | Dual-task Level | -792 | -664 | 128 | 3,76510123 | 0,00659868 | 0,434756442 | beta |
| Sensorimotor | Cognitively_Enhanced_DT vs Motor_Enhanced_DT | Dual-task Level | -92 | -16 | 76 | 3,668083535 | 0,015296941 | 0,423553803 | beta |
| Sensorimotor | Cognitively_Enhanced_DT vs Cognitive-Motor_Enhanced_DT | Dual-task Level | -792 | -664 | 128 | 4,230263787 | 0,0014997 | 0,488468787 | beta |
| Sensorimotor | Easy_DT vs Motor_Enhanced_DT | Dual-task Level | 36 | 632 | 596 | 5,843306102 | 0,004874025 | 0,67472687 | beta |
| Sensorimotor | Easy_DT vs Cognitive-Motor_Enhanced_DT | Dual-task Level | 36 | 632 | 596 | 6,081765438 | 0,006007132 | 0,702261783 | beta |
| Sensorimotor | Cognitively_Enhanced_DT vs Motor_Enhanced_DT | Dual-task Level | 12 | 660 | 648 | 5,469514143 | 0,002191869 | 0,631565093 | beta |
| Sensorimotor | Cognitively_Enhanced_DT vs Cognitive-Motor_Enhanced_DT | Dual-task Level | 12 | 660 | 648 | 5,653332845 | 0,003045545 | 0,652790648 | beta |

**Supplementary methods**

We conducted supplementary analyses on ray time frequency power using the same model as in the manuscript. The design matrix was constructed to model the factorial structure of the experiment, including body visualization scenarios, dual-task level, and the condition-by-dual-task interaction and participant as random intercept. Time-frequency power in the following bands (alpha, beta, theta) and ROIs (frontocentral and sensorimotor), from 800 ms before to 800 ms after the event. The design matrix was permuted 5,000 times to estimate a null distribution, and main effects and interaction effects were first tested, followed by pairwise comparisons.

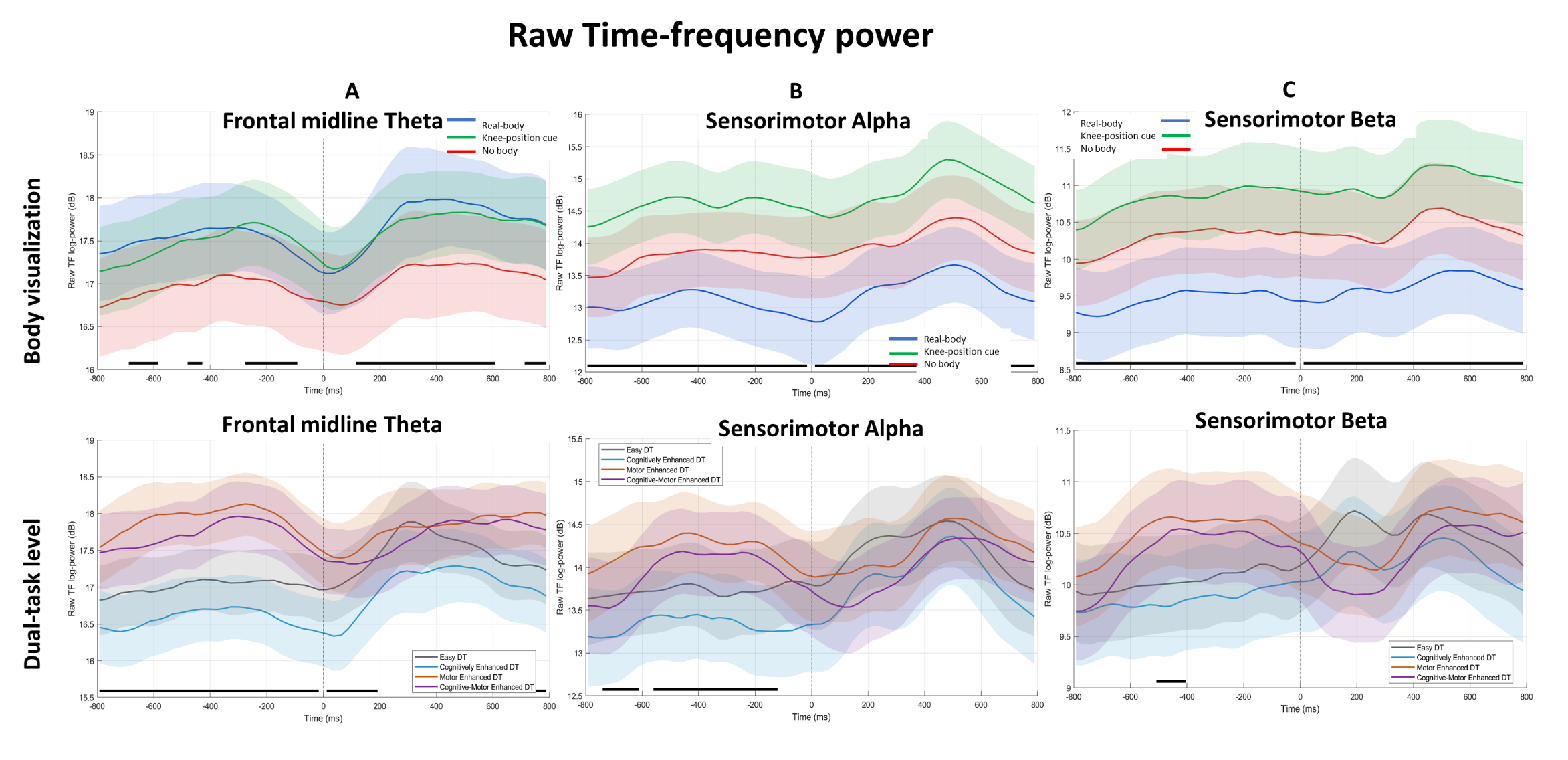

Figure S1. **Raw time frequency across body visualization conditions and dual-task levels in (A) frontal midline theta, (B) sensorimotor alpha, and (C) sensorimotor beta bands**. Panels show raw power in decibels with corresponding standard errors of the mean (SEM). **Time zero represents the moment of clearing the obstacles during the stepping execution**. Solid black lines at the bottom indicate time windows with significant pairwise differences identified by cluster-based permutation testing. For baseline (-300 -100 ms) effects of body visualization, (A) theta power was higher during visualization of knee-position dans during the real-body and no-body visualization condition. (B-C) In the alpha and beta band, no-body visualization and that of knee-position cue showed more power than real-body visualization with knee-position cue also differing from no-body. The tonic effect of body visualization is visible in all bands (A-C) with modulation throughout the epoch. For dual-task level, pre-event effects were observed across all bands (A-B): the motor-enhanced and cognitive–motor-enhanced levels showed comparable power and both exhibited more power than the easy and cognitive-enhanced levels.
